# Systemic Inflammation Mediates the Relationship between Obesity and Health Related Quality of Life

**DOI:** 10.1101/231720

**Authors:** Jeffrey Wilkins, Palash Ghosh, Juan Vivar, Bibhas Chakraborty, Sujoy Ghosh

**Affiliations:** Biomedical Biotechnology Research Institute, North Carolina Central University, 1801 Fayetteville Street, NC 27707, USA; Centre for Quantitative Medicine, Duke-NUS Medical School, 8 College Road, Singapore 169857; Center for Tobacco Products, Food and Drug Administration, 10903 New Hampshire Avenue, Silver Spring, MD 20993, USA; Program in Cardiovascular & Metabolic Disorders & Centre for Computational Biology, Duke-NUS Medical School, 8 College Road, Singapore 169857

**Keywords:** Obesity, inflammation, healthy days, health-related quality of life, mediation

## Abstract

**Background:** At the population level, obesity has been reported to be positively associated with low-level chronic inflammation, and negatively associated with several indices of health-related quality of life (HRQOL). It is however not clear if obesity-associated inflammation is partly responsible for the observed negative associations between obesity and HRQOL. The present study investigates this question by testing the hypothesis that systemic inflammation is a mediator of the observed association between obesity and a specific HRQOL index called “healthy days”, as measured via a subset of the CDC HRQOL-4 questionnaire.

**Methods:** Demographic, body mass index (BMI), C-reactive protein (CRP), inflammatory disease status, medication use, smoking, and HRQOL data were obtained from NHANES (2005-2008) and analyzed using sampling-weighted generalized linear models. Both main effects and interaction effects were analyzed to evaluate possible mediator-outcome confounding. Model robustness was tested via sensitivity analysis. Prior to model development, data was subjected to multiple imputation in order to mitigate information loss from survey non-response. Averaged results from the imputed datasets were reported in the form of odds ratios (OR) and confidence intervals (CI).

**Results:** Obesity (BMI >30kg/m^2^) was positively associated with poor physical healthy days (OR: 1.59, 95% CI: 1.15-2.21) in unadjusted models. ‘Elevated’ and ‘clinically raised’ levels of the inflammation marker CRP were also positively associated with poor physical healthy days (OR= 1.61, 95% CI: 1.23-2.12, and OR= 2.45, 95% CI: 1.84-3.26, respectively); additionally ‘clinically raised’ CRP was positively associated with mental unhealthy days (OR= 1.66, 95% CI: 1.26-2.19). The association between obesity and physical HRQOL was rendered non-significant in models including CRP. Association between ‘elevated’ and ‘clinically raised’ CRP and physical unhealthy days remained significant even after adjustment for obesity or inflammation-modulating covariates (OR= 1.36, 95% CI :1.02-1.82, and OR= 1.75, 95% CI: 1.21-2.54, respectively).

**Conclusions:** Systemic inflammation is a significant mediator of the association between obesity and physical unhealthy days. and is also an independent determinant of physical and mental unhealthy days. Importantly, elevated inflammation below the clinical threshold is also negatively associated with physical healthy days and may warrant more attention from a population health perspective than currently appreciated.

## BACKGROUND

Inflammation is a necessary component of host defenses against infections and injury, but can also contribute to the pathophysiology of several chronic diseases [1, 2]. Chronic systemic inflammation provides a unifying pathological mechanism for many seemingly unrelated diseases including arthritis, asthma, atherosclerosis, diabetes, congestive heart failure, Crohn’s disease, Alzheimer’s disease and several forms of cancers [3]. Extensive research also points to a strong association between inflammation and obesity, a global epidemic and clear public health concern [4-6]. Obesity (body mass index, BMI≥30kg/m^2^) is often co-extant with low-level chronic inflammation that is reflected in elevated levels of circulating CRP [7, 8]. Obesity-associated inflammation leads to insulin resistance, endothelial dysfunction, and eventually, a significant elevation of cardio-metabolic disease risk [9, 10].

From a population health perspective, effective interventions and optimized predictions for future health-care costs require a better quantitative understanding of chronic conditions such as inflammation and obesity and their relationship to indices of public health. One index that captures the population level effects of chronic conditions is the health related quality of life (HRQOL) [11]. HRQOL, a self-reported measure of physical and mental functioning and well-being, is increasingly used to assess the effects of chronic illness, treatments, and short- and long-term disabilities.

Previous studies have generally demonstrated a negative correlation between excess adiposity and HRQOL in different populations [12-16]. Much less is known, however, on the effects of chronic inflammation on HRQOL, and whether inflammation can mediate the association between obesity and HRQOL. Studies investigating inflammation and HRQOL have either focused on small cohorts targeting specific inflammatory disorders [17-19] or interrogated inflammation and HRQOL as separate endpoints. Here we perform a mediation analysis, including potential mediator-outcome confounder analysis [20], to examine the relationship between systemic inflammation, obesity and the HRQOL measures of physical and mental healthy days (based on the CDC HRQOL-4 questionnaire), from survey data derived from the National Health and Nutrition Examination Survey, (NHANES) 2005-2008. Our findings point to new insights involving obesity, inflammation and HRQOL, with potentially important implications for public health.

## METHODS

### Study design and Participants

For this cross-sectional study, data was downloaded from the National Health and Nutrition Examination Survey (NHANES) collection (years 2005 through 2008, http://www.cdc.gov/nchs/nhanes/nhanes_questionnaires.htm). NHANES is conducted by the Center for Disease Control’s National Center for Health Statistics (CDC-NCHS) to assess the health and nutritional status of a representative civilian, non-institutionalized US population using a multistage, stratified, clustered probability design [21]. Data for the variables of interest were available for a total of 6325 adults (aged 20-75 years, BMI≥18.5 kg/m^2^) and included missing values. Data collected included demographic information, health-related questionnaire, and laboratory data on C-reactive protein (CRP, a marker of systemic inflammation). Data for confounding factors that can influence inflammation status and HRQOL outcomes such as relevant medical conditions (asthma, arthritis, heart disease, and cancer), anti-inflammatory/analgesic medication, and lifestyle choice (smoking) were also downloaded for statistical modeling. To account for the complexity of survey design including oversampling, survey non-response, or post-stratification issues, NHANES assigns sample ‘weights’ were also downloaded (additional details available from https://www.cdc.gov/nchs/tutorials/nhanes/SurveyDesign/Weighting/OverviewKey.htm, and in **Supplementary Text 1**).

### Health related quality of life (HRQOL) measures

Quality of life was assessed by using a subset of the CDC HRQOL-4 questionnaire, developed to assess physical and mental health in the general U.S population [22, 23]. The HRQOL-4 questionnaire **(Supplementary Text 2)** uses self-reported measures of healthy and unhealthy days as indicators of HRQOL, and have undergone cognitive testing and criterion validity with the Short-Form 36, content and construct validity, predictive validity, internal consistency, test-retest reliability, and measurement invariance in persons with and without disability [24, 25]. The core Healthy Days consists of four questions focusing on the participant’s general health status (Question-1), number of physical unhealthy days in the 30 days preceding the survey (Question-2), number of mental unhealthy days in the 30 days preceding the survey (Question-3), and number of days with activity limitations in the 30 days preceding the survey (Question-4). Question-1 is a predictor of mortality and chronic disease conditions [26], questions −2 and −3 assess recent physical symptoms and mental or emotional distress, respectively, and question-4 measures perceived disability and lost productivity [23]. Only responses to survey Questions-2 and -3 were used in the current study since the focus of the analysis was to determine effects specifically on physical and mental health. Throughout this analysis, an *increase* in the number of physical/mental unhealthy days has been used to indicate poorer health outcomes.

### Coding of variables

Participants were divided into 3 categories by age (class 1, 20-44 yrs.; class 2, 45-65 yrs.; class 3 >65 yrs.) and categorized by race/ethnicity as Mexican-American (1), Other Hispanic (2), Non-Hispanic White (3), Non-Hispanic Black (4), and Other (5). Obesity was measured by body mass index (BMI) based on self-reported weight in kilograms divided by measured height in meters-square. Respondents were broadly categorized into 3 BMI groups: normal weight (BMI 18.5-24.9), overweight (BMI 25-29.9), and obese (BMI≥30). Systemic inflammation was measured via blood CRP levels (mg/dl) and grouped into 3 classes according to Visser et al. [8] – Class 1(‘non-elevated CRP’, <0.22mg/dl), Class 2 (‘elevated CRP’, ≥0.22 and <1.0 mg/dl) and Class 3 (‘clinically raised CRP’; ≥1.0 mg/dl), respectively. Each medical condition, including asthma (MCQ010), arthritis (MCQ160A), cancer (MCQ220), or any heart disease (a composite variable derived from a positive diagnosis of any one of congestive heart failure (MCQ160B), coronary heart disease (MCQ160C), or heart attack (MCQ160E)) were dichotomized into ‘0’ and ‘1’ categories where ‘1’ indicates a positive response to the question of whether there ever was a diagnosis of the relevant condition by a doctor or healthcare professional. Smoking status was also dichotomized, with ‘1’ assigned to individuals who are current smokers. The use of common analgesic and anti-inflammatory medications (aspirin, acetaminophen, ibuprofen and naproxen) was also recorded with ‘1’ indicating use of drug and ‘0’ otherwise. Although acetaminophen is more widely prescribed as an analgesic and antipyretic, previous studies have reported effectiveness of acetaminophen against lower grade inflammation [27] and acetaminophen overdose has also been shown to reduce circulating CRP levels [28]. Based on these findings, and the close association between inflammation and physical pain, we included the use of acetaminophen in the analysis. For logistic regression analysis, each outcome variable (HRQOL measures) was dichotomized into ≤15 or >15 days of poor physical (HSQ470) or mental health (HSQ480) with >15 unhealthy days denoted by 1, and 0 otherwise.

### Statistical Analysis

All statistical analysis was conducted using SAS, version 9.1 (SAS Institute Inc., Cary, NC, USA) or R (version x64 3.2.3). Data was analyzed using sampling weighted generalized linear models. Both unadjusted and adjusted models linking BMI and CRP to the outcome variables (HSQ470 and HSQ480) were constructed, with adjustments for possible inflammation-regulating medical conditions (asthma, arthritis, heart disease, and cancer), use of over the counter anti-inflammatory/pain medications (acetaminophen, aspirin, ibuprofen and naproxen), and lifestyle (current cigarette smoking status). As several of the surveyed variables contained missing values, any attempt to analyze only complete cases severely reduced the total number of observations, leading to potentially biased estimates. To circumvent this problem, we used multiple imputation [29] to estimate probable value ranges for incomplete observations. The original coding for the missing values included bona-fide missing values, plus other types of non-response such as “don’t know" (1517 total instances), and “refused to answer” (2 total instances). We converted all missing and non-response cases into missing values, as recommended by the NHANES analytical guidelines [30]. Five imputed datasets were generated according to multiple imputation procedures described by Rubin [29]. Each of these “completed’ datasets were individually analyzed using sampling weighted generalized linear models (GLM), via the *survey* package in R [31]. For each of the imputed datasets, *m* = 1…5, we obtained the estimate of regression coefficient as *β_m_* along with the standard error *s_m_*. The overall estimate was obtained by averaging the individual estimates from the imputed datasets as

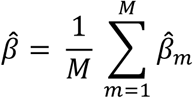

The estimated variance for 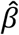 is given by

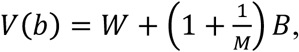

where 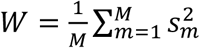, and 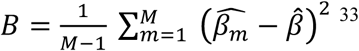.

Due to the difficulties in combining and interpreting the combined p-values arising from the above analysis, we have chosen to report results in terms of the estimated odds ratio 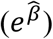 and their 95% confidence interval for all analysis. The odds ratios (OR) were obtained by exponentiation of the average regression coefficient, 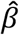.

## RESULTS

### Participants

Data for subjects 18-20 years of age were excluded due to the large excess of missing values in this group and to prevent complications from differential growth patterns in childhood and adolescence where the usual BMI categories do not apply [32]. The general characteristics of the survey respondents are listed in **Table 1**. The average age of the sampled population was 51.3 years (±17.85 years), with approximately 49% male subjects. The mean physical and mental unhealthy days reported was 4.49(±8.71) and 4.09 (±8.28) days, respectively. The average BMI was 29.34(±6.77) kg/m^2^, with approximately 27% normal-weight, 35% overweight, and 38% obese subjects. The mean CRP level was 0.46(±0.89) mg/dl, with approximately 51% ‘normal’, 37% ‘elevated’ and 11% ‘clinically raised’ categories. Approximately 22% of the subjects were current smokers (SMQ040), whereas nearly 14% had a diagnosis of asthma (MCQ010). Similar medical diagnosis for arthritis (MCQ160A) and cancer/malignancy (MCQ220) were 32% and 10%, respectively. Approximately 9.5% of the population was positive for “any heart disease”. Finally, nearly 14% of the subjects reported using one or more of the analgesic/ anti-inflammatory medications.

**Table 1.**
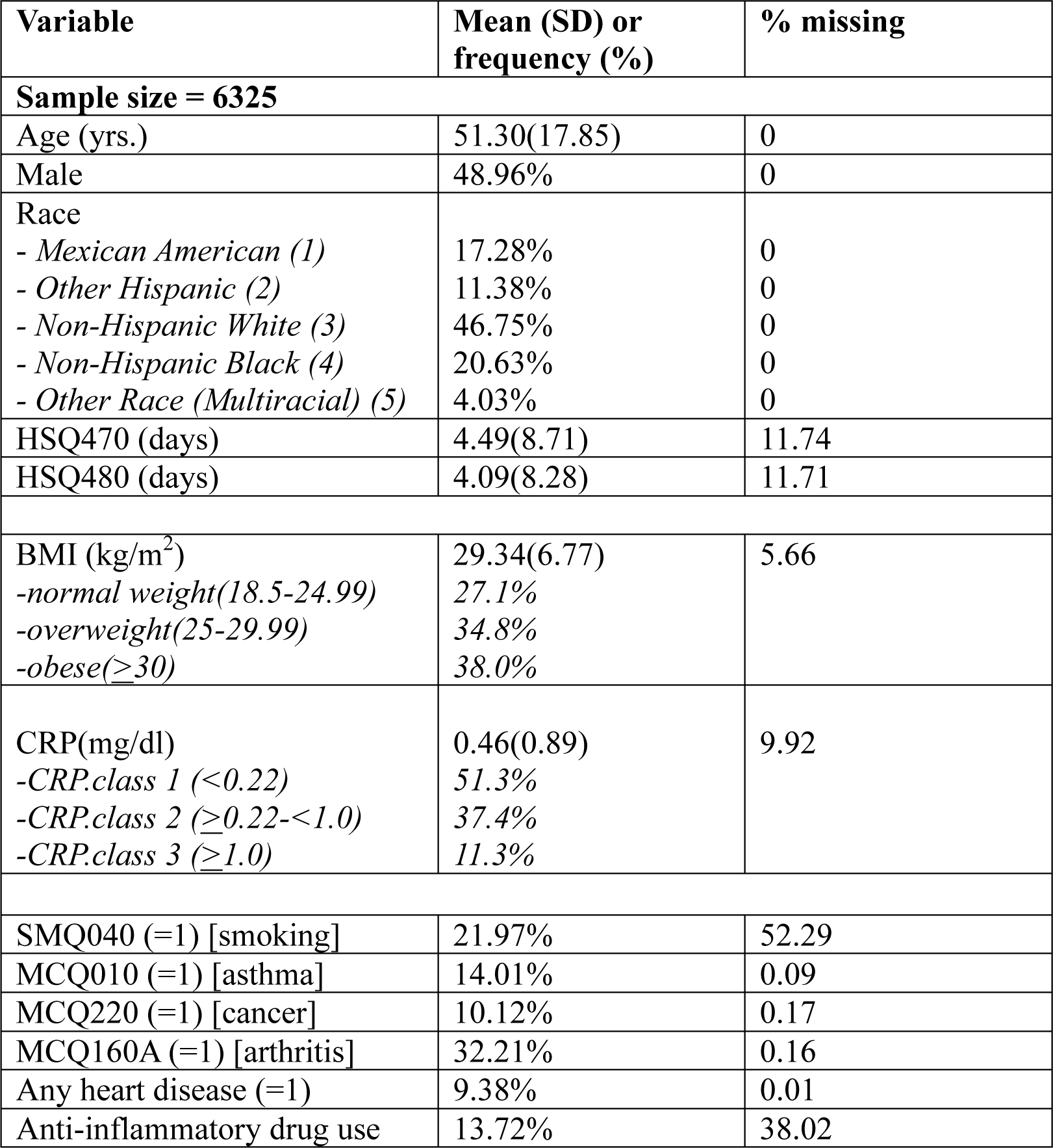
***Demographic and Medical Characteristics of Study Subjects.** Data is presented as mean(SD) for continuous variables and as frequency(%) for categorical variables. The percent of data missing for each variable is indicated. Inflammation-related variables are coded (as per NHANES 2005-2008) as follows: SMQ040 (current smoking status), MCQ010 (medical diagnosis of asthma), MCQ220 (medical diagnosis of cancer), MCQ160A (medical diagnosis of arthritis), Any heart disease (medical diagnosis of one or more of heart attack, congestive heart failure or coronary heart disease)*

| Variable | Mean (SD) or frequency (%) | % missing |
| --- | --- | --- |
| **Sample size = 6325** |  |  |
| Age (yrs.) | 51.30(17.85) | 0 |
| Male | 48.96% | 0 |
| Race |  |  |
| - Mexican American (1) | 17.28% | 0 |
| - Other Hispanic (2) | 11.38% | 0 |
| - Non-Hispanic White (3) | 46.75% | 0 |
| - Non-Hispanic Black (4) | 20.63% | 0 |
| - Other Race (Multiracial) (5) | 4.03% | 0 |
| HSQ470 (days) | 4.49(8.71) | 11.74 |
| HSQ480 (days) | 4.09(8.28) | 11.71 |
| BMI (kg/m<sup>2</sup>) | 29.34(6.77) | 5.66 |
| -normal weight(18.5-24.99) | 27.1% |  |
| -overweight(25-29.99) | 34.8% |  |
| -obese($\geq 30$) | 38.0% | |
| CRP(mg/dl) | 0.46(0.89) | 9.92 |
| -CRP.class 1 (<0.22) | 51.3% |  |
| -CRP.class 2 ($\geq 0.22$-<1.0) | 37.4% | |
| -CRP.class 3 ($\geq 1.0$) | 11.3% | |
| SMQ040 (=1) [smoking] | 21.97% | 52.29 |
| MCQ010 (=1) [asthma] | 14.01% | 0.09 |
| MCQ220 (=1) [cancer] | 10.12% | 0.17 |
| MCQ160A (=1) [arthritis] | 32.21% | 0.16 |
| Any heart disease (=1) | 9.38% | 0.01 |
| Anti-inflammatory drug use | 13.72% | 38.02 |

### Relationship between body mass index and C-reactive protein

Quantile-quantile plots demonstrated that CRP and BMI values were better approximated to the normal distribution after log transformation (data not shown). We carried out linear regression to determine the association between CRP levels and BMI. Taking log CRP as the dependent variable and log BMI as the predictor variable, the regression coefficient of BMI was 2.69 (95%CI: 2.50, 2.88) (**Supplementary Table 1**), indicating a statistically significant association between BMI and CRP in the study population.

### Relationship of body mass index (BMI) and C-reactive protein (CRP) to physical and mental healthy days

Considering physical unhealthy days (HSQ470) as a binary response, we performed logistic regression with BMI groups (normal, overweight and obese) (normal group as reference) (**Table 2, model 1**). The estimated odds ratio for overweight subjects was 1.06 (95% CI: 0.76, 1.47) and that for obese subjects was 1.59 (95% CI: 1.15, 2.21). Thus, compared to a normal-weight person, an overweight person (25 ≤ BMI ≤ 29.9) was 1.06 times more likely, and an obese person (BMI ≥30) 1.59 times more likely to experience >15 physical unhealthy days in a month. Only the estimated OR for the obese, but not overweight, individuals were statistically significant (95% CI excluded 1). In contrast, neither overweight nor obese individuals were significantly associated to mental unhealthy days (95% CI includes 1) (**Table 2, model 2**).

**Table 2.** ***Relationship of Physical and Mental Healthy Days to BMI and CRP levels.** Results include estimates of odds ratio (OR) and corresponding 95% confidence intervals under different models indexed by varying dependent variables. The OR is interpreted as the relative changes in odds for physical (HSQ470>15 days) or mental (HSQ480>15 days) unhealthy days upon changes in the categories of the explanatory variables (BMI and/or CRP).*

| Model | Dependent Variable | Parameter | OR (95% CI) |
| --- | --- | --- | --- |
| Model 1 | HSQ470 | (Intercept) | 0.06 (0.04,0.09) |
|  |  | overweight | 1.06 (0.76,1.47) |
|  |  | obese | 1.59 (1.15,2.21) |
| Model 2 | HSQ480 | (Intercept) | 0.06 (0.05,0.09) |
|  |  | overweight | 1.20 (0.86,1.68) |
|  |  | obese | 1.25 (0.89,1.75) |
| Model 3 | HSQ470 | (Intercept) | 0.06 (0.04,0.07) |
|  |  | CRP.class (2) | 1.61 (1.23,2.12) |
|  |  | CRP.class (3) | 2.45 (1.84,3.26) |
| Model 4 | HSQ480 | (Intercept) | 0.07 (0.05,0.09) |
|  |  | CRP.class (2) | 1.05 (0.79,1.40) |
|  |  | CRP.class (3) | 1.66 (1.26,2.19) |
| Model 5 | HSQ470 | (Intercept) | 0.05 (0.04,0.07) |
|  |  | overweight | 0.98 (0.70,1.35) |
|  |  | obese | 1.26 (0.92,1.74) |
|  |  | CRP.class (2) | 1.51 (1.14,2.0) |
|  |  | CRP.class (3) | 2.21(1.55,3.16) |

To further assess the relationship between the obesity class and physical/mental unhealthy days, we focused only on obese subjects (BMI≥30), divided into 5 subclasses according to increasing values of BMI (**Supplementary Tables 2 and 3**). Subjects in the two highest classes of obesity (class IV, BMI 50.0-59.9 and class V, BMI ≥60) were found to be significantly associated to physical unhealthy days, compared to baseline (class I obesity, BMI 30.0-34.9). Only class V obesity subgroup was found to be significantly associated to HSQ480, with higher BMI associated with a reduced probability for mental unhealthy days. Although this finding is counterintuitive, we note that the statistical estimates may be unstable due to the very low subject numbers in this group (15 individuals, < 1% of total BMI≥30 population).

Next, we assessed the relationship between plasma CRP levels (with CRP.Class 1 as reference) and the number of physical unhealthy days (**Table 2,model 3**). The estimated OR of CRP.Class (2) was 1.61 (95% CI: 1.23, 2.12) and that of CRP.Class (3) was 2.45 (95% CI: 1.84, 3.26), suggesting statistically significant associations for both CRP categories. The association between CRP.Class(2) to mental unhealthy days (HSQ480) was not significant (OR=1.05, 95% CI: 0.79, 1.40); however, CRP.Class(3) was significantly associated to mental unhealthy days, (OR=1.66, 95% CI: 1.26, 2.19) **(Table 2, model 4)**.

We next modeled both BMI groups and CRP.Class as explanatory variables to ascertain their relative contribution to physical unhealthy days. The estimated odds ratios were 0.98 and 1.26 for overweight and obese BMI groups, respectively, and, 1.51 and 2.21 for CRP.Class(2) and CRP.Class(3), respectively (**Table 2, model 5**). However, both the 95% CIs corresponding to the overweight (95% CI: 0.70, 1.35) and obese group (95% CI: 0.92, 1.74) now included 1, whereas the corresponding CIs for CRP.Class(2) (95% CI: 1.14, 2.00) and CRP.Class(3) (95% CI: 1.55, 3.16) excluded 1. In other words, when both CRP and BMI are included as explanatory variables in the same model, the significant associations observed earlier between BMI level and physical unhealthy days was no longer present, suggesting systemic inflammation as mediating the effect **(Figure 1)**.

**Figure 1.**
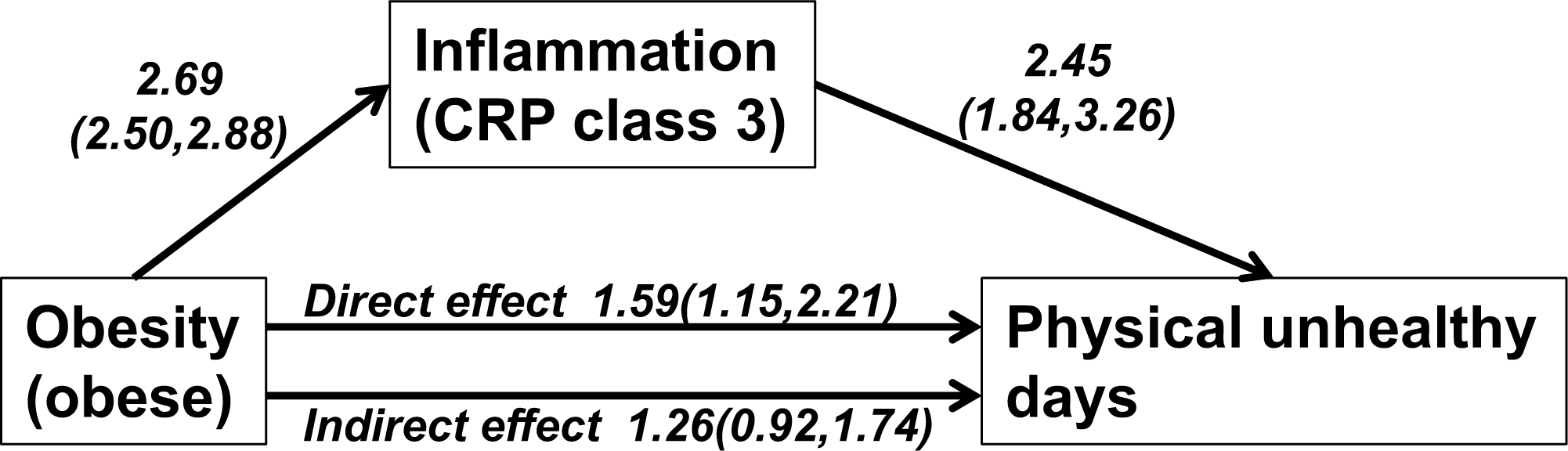
Effect of adjusting for inflammation on the estimate of obesity-physical unhealthy days association in NHANES subjects. The ‘direct effect’ estimates the odds ratio (and confidence intervals) for the association of adult obese subjects to the number of physical unhealthy days, a measure of HRQOL. The ‘indirect effect’ estimates the same association following the inclusion of clinically raised systemic inflammation (measured as C-reactive protein) as a possible mediator. The direct associations of obesity to inflammation and of inflammation to physical unhealthy days are also shown.

### Effect modification analysis

We carried out an effect modification analysis on the relationship of HSQ470 with BMI and CRP by including gender, age-class and race in the models. The interaction effects between 'overweight and gender’ and 'obese and gender’ were significant (**Table 3**). For example, within the overweight category, a male was 0.42 times less likely to experience >15 physical unhealthy days compared to a female. All other interactions were non-significant. For HSQ480 (mental unhealthy days), we observed significant interaction effects due to ‘obese and gender’; ‘CRP.class(3) and gender’; ‘overweight and Ageclass(2)’; ‘obese and Ageclass(2)’; ‘CRP.class(3) and Ageclass(2)’; ‘CRP.class(3) and Ageclass(3)’, and, ‘overweight and Race-5’ (**Supplementary Table 4**). All other interactions effects were non-significant. These results suggest that the observed association between adiposity or CRP and physical/mental healthy days are modifiable to some extent by age and gender. The apparent modification of the association between overweight and HSQ480 by Race-5 has to be interpreted with caution due to the very low numbers of subjects belonging to this category (<5% of the surveyed population, **Table 1**).

**Table 3:**
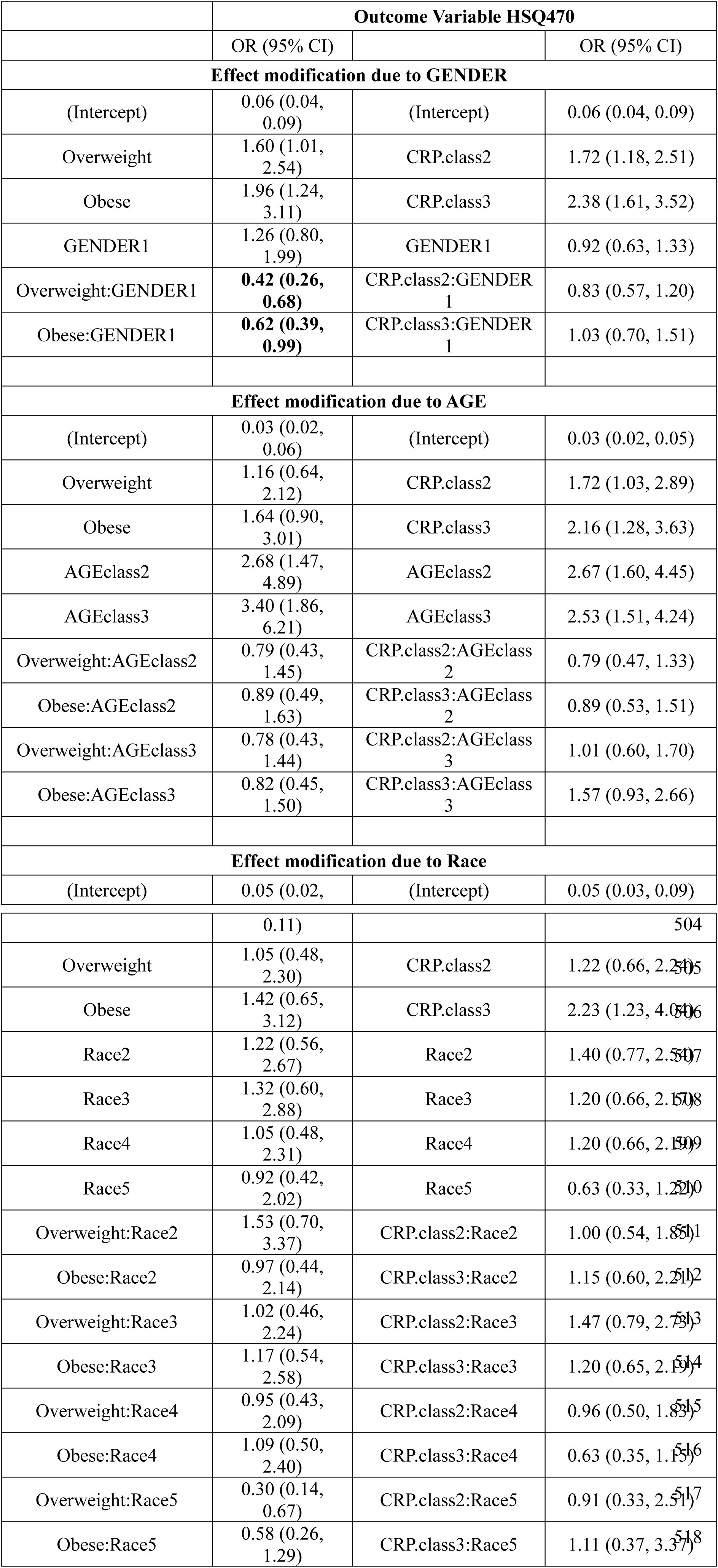
***Effect modification for outcome variable HSQ470.** The modification of the association between physical healthy days (HSQ470) and BMI or CRP was investigated. Data was analyzed using sampling weighted generalized linear models (logistic) as described under Methods.*

### Sensitivity analysis

We performed sensitivity analysis on the relationship of HSQ470 to BMI groups and CRP classes respectively, by varying the cut-off value for HSQ470=1 from 15 to 12, 13, 14, 16, 17 and 18 days. The obese class was significantly associated to HSQ470 for all the cut-off values tested with relatively stable odds ratio estimates **(Table 4).** On the other hand, the odds-ratios for overweight were non-significant for all HSQ470 cut-off values tested, consistent with the original findings. Similarly, the odds-ratios corresponding to the different CRP classes (2 and 3) with different HSQ470 cut-off values were significant, agreeing again with the primary results (HSQ470 cut-off value=15). These results suggest that the identified associations between BMI or CRP and HRQOL are robust to the threshold used for defining physical unhealthy days.

**Table 4:** ***Sensitivity Analysis with respect to different cut-off values of HSQ470 vs BMI and CRP.** The threshold for physical unhealthy days was varied from 12-18 days and the effects on the association to BMI or CRP classes was evaluated (upper and lower panels of table, respectively). Data was analyzed by sampling weighted generalized linear models as described under Methods.*

|  | OR (95% CI) (outcome variable HSQ470 vs BMI Class) |  |  |  |  |  |
| --- | --- | --- | --- | --- | --- | --- |
| Cut-off of HSQ470 | 12 | 13 | 14 | 16 | 17 | 18 |
| Intercept | 0.08<br>(0.06,0.11) | 0.08<br>(0.06,0.11) | 0.07<br>(0.05,0.1) | 0.06<br>(0.04,0.08) | 0.06<br>(0.04,0.08) | 0.06<br>(0.04,0.08) |
| Overweight | 1.21<br>(0.91,1.59) | 1.19<br>(0.9,1.58) | 1.2<br>(0.89,1.61) | 1.03<br>(0.74,1.43) | 1.02<br>(0.73,1.42) | 1.02<br>(0.73,1.42) |
| Obese | **1.69**<br>**(1.27,2.23)** | **1.7**<br>**(1.28,2.25)** | **1.66**<br>**(1.23,2.23)** | **1.6**<br>**(1.15,2.23)** | **1.58**<br>**(1.13,2.2)** | **1.54**<br>**(1.1,2.15)** |
|  | OR (95% CI) (outcome variable HSQ470 vs CRP Class) |  |  |  |  |  |
| Cut-off of HSQ470 | 12 | 13 | 14 | 16 | 17 | 18 |
| Intercept | 0.08<br>(0.06,0.1) | 0.08<br>(0.06,0.1) | 0.07<br>(0.05,0.09) | 0.05<br>(0.04,0.07) | 0.05<br>(0.04,0.07) | 0.05<br>(0.04,0.07) |
| CRP.Class2 | **1.54**<br>**(1.22,1.96)** | **1.53**<br>**(1.2,1.93)** | **1.55**<br>**(1.21,1.98)** | **1.7**<br>**(1.29,2.24)** | **1.7**<br>**(1.28,2.24)** | **1.73**<br>**(1.3,2.28)** |
| CRP.Class3 | **2.34**<br>**(1.83,2.99)** | **2.36**<br>**(1.84,3.03)** | **2.46**<br>**(1.89,3.2)** | **2.59**<br>**(1.94,3.45)** | **2.64**<br>**(1.97,3.54)** | **2.7**<br>**(2.01,3.62)** |

### Relationship of CRP to physical and mental unhealthy days after adjustment for other sources of inflammation

We next investigated whether the effects of CRP classes on physical unhealthy days could be confounded by some of the most common sources of inflammation encountered in the study population (mediator outcome confounding). We carried out multivariable logistic regression analysis by including demographics (age, gender), pro-inflammatory medical conditions, use of common anti-inflammatory/pain medications, and current smoking status, in addition to CRP and BMI categories in the model (**Table 5**). The CRP.Class variable remained significantly associated to physical unhealthy days, for both the CRP.Class (2) (OR=1.36, 95% CI:1.02, 1.82) and CRP.Class (3) (OR=1.75, 95% CI:1.21, 2.54), even after adjustment. A similar analysis against mental unhealthy days showed the association of CRP classes and BMI groups to be non-significant, although significant associations were observed for presence of asthma, presence of arthritis, current smoking status, occurrence of any heart disease and gender **(Supplementary Table 5)**.

**Table 5:** ***Multivariable logistic regression analysis of the association of CRP to physical unhealthy days.** Results include estimates of odds ratio (OR) and corresponding 95% confidence intervals. The OR is interpreted as the increase in odds for physical (HSQ470>15 days) unhealthy days upon changes in the categories of the explanatory variables. Data was analyzed using sampling weighted generalized linear models (logistic) as described under Methods.*

| Parameter | OR (95% CI) |
| --- | --- |
| (Intercept) | 0.02 (0.01, 0.04) |
| Overweight | 0.93 (0.66, 1.30) |
| Obese | 1.08 (0.76, 1.52) |
| CRP.class (2) | 1.36 (1.02, 1.82) |
| CRP.class (3) | 1.75 (1.21, 2.54) |
| Anti-inflammatory Drug Use (1) | 2.42 (1.76, 3.35) |
| AGEclass (2) | 1.83 (1.31, 2.57) |
| AGEclass (3) | 1.67 (1.11, 2.50) |
| MCQ010 (1) | 1.37 (1.01, 1.85) |
| MCQ220 (1) | 1.36 (0.95, 1.94) |
| MCQ160A (1) | 2.33 (1.72, 3.14) |
| GENDER (1) | 0.89 (0.69, 1.16) |
| SMQ040 (1) | 1.16 (0.84, 1.61) |
| Any Heart Disease (1) | 1.65 (1.15, 2.35) |

## DISCUSSION

The present study was undertaken to better define the relationship between obesity, systemic inflammation and measures of HRQOL. We used data from a US population based survey (NHANES 2005-2008) to estimate effects of increasing body mass and increasing inflammation on the number of physical and mental unhealthy days reported by participants, in a mediation framework **(Figure 1)**. We also tested the impact of common inflammation regulators (inflammatory disease, anti-inflammatory drug use, and smoking) on the association between the inflammation marker CRP, and HRQOL (based on the CDC HRQOL-4 questionnaire). Compared to the more detailed Medical Outcomes Study Short Form 36 (SF-36), the CDC’s ‘‘healthy days’’ serves as a simple proxy measure of HRQOL. It measures perceptions of physical and mental health using one question each, eliminating the need for complex weighting factors to calculate summary scores.

In previous studies, Hassan et al. [13] assessed a US-based sample with the CDC-HRQOL-4 and reported greater likelihood of poor physical and mental quality of life in participants with obesity. Renzaho et al. [15] sampled an Australian population with SF-36 and found that physical, but not mental, QoL scores were negatively associated with BMI. Serrano-Aguilar et al. [16] analyzed a European sample using the EuroQol-5d assessment and found that participants with BMI≥40 had lower HRQOL scores than normal weight participants. These findings agree with the positive association between BMI and number of physical unhealthy days observed in the current study in unadjusted models, and also support the lack of association between BMI and mental unhealthy days[15]. Importantly however, the association between BMI and physical HRQOL was non-significant after systemic inflammation (CRP levels) was included as a mediator **(Table 2, model 5)**, or other variables that contribute towards such inflammation **(Table 5)**. A summary of the major findings s presented via the directed acyclic graph [33] in **Figure 1.** We speculate that the observed earlier associations between BMI and HRQOL could be mediated by the chronic inflammation that co-exists in obese subjects. However, methodological differences between the current study and previously published reports should also be noted. While our study used the number of healthy days as the QoL metric, previous studies utilized composite HRQOL measures, based on a summation over several sub-domain scores. Additionally, differences in the sampled populations between the studies could also potentially influence the current findings. The cross-sectional design of the current study further limits the ability to infer causal relationships, e.g. one cannot distinguish if inflammation causally affected HRQOL, or if HRQOL was affected by some other factors that also led to inflammation. Finally, since the assessment by CDC HRQOL-4 is based on self-reporting, the study results are also potentially susceptible to the risk of recall error and misreport.

A second significant finding from the current study is that systemic CRP levels are positively and significantly associated with the number of physical and mental, unhealthy days, even after adjustments for sex, age, pro-inflammatory co-morbidities, and anti-inflammatory/analgesic drug use. The observation that even sub-clinical CRP levels affect HRQoL may have important consequences for public health. Visser et al. [8] introduced the classification scheme of sub-clinical ‘elevated CRP (≥0.22 mg/dl)’ and ‘clinically raised CRP (≥1.0 mg/dl)’ and identified an association of the former with overweight and obesity. In other studies, sub-clinical CRP has been associated with increased risk of cardiovascular disease-related mortality in healthy subjects [34].

## CONCLUSION

In conclusion, a population-based mediation analysis investigating the roles of obesity and systemic inflammation on indices of health-related quality of life suggests inflammation as a strong mediator of the negative associations between body mass index and the number of reported physical healthy days. Our findings further suggest that sub-clinical inflammation is possibly also an independent predictor of quality of life domains in the general population. In light of these observations, the relationship of systemic inflammation to patient’s health may need to be re-assessed and a distinction made between very high CRP levels due to acute events, and lower (but possibly chronic) elevations in CRP that can still significantly affect health related quality of life, and therefore should be targeted for clinical management.

## ABBREVIATIONS

BMI: Body Mass Index
CRP: C-reactive protein
HRQOL: Health Related Quality of Life
HSQ470: Health Status Questionnaire 470
HSQ480: Health Status Questionnaire480
NHANES: National Health and Nutrition Examination Survey

## DECLARATIONS

### Ethics approval and consent to participate

The sample in the current study included adults aged 20 to 75 years, with BMI≥18.5 kg/m^2^, who completed the examination component in 2005–2006 or 2007–2008. The NHANES surveys are subject to CDC-NCHS Ethics Review Board to ensure appropriate human subject protections, in compliance with 45 Code of Federal Regulations, part 46 [35].

### Consent for publication

Not applicable

### Availability of data and materials

The datasets used and/or analyzed during the current study are available from the corresponding author on reasonable request. The original NHANES data is freely available from https://www.cdc.gov/nchs/nhanes/index.htm

### Competing interests

The authors declare that they have no competing interests.

### Funding

This work was supported by a NCCU-BBRI fellowship to JW and funding from the National Institutes of Health (HL059868, MD000175 and DK088319), American Heart Association (AHA10SDG4230068) National Medical Research Council, Ministry of Health, Singapore (WBS R913200076263) to SG; funding from the Duke-NUS Medical School, Singapore, to BC.

### Author contributions

**JW** (acquisition and interpretation of data, writing of manuscript); **PG** (analysis of data, writing of manuscript); **JV** (analysis of data); **BC** (interpretation of data, review of statistical analysis); **SG** (conception and supervision of study, interpretation of data, writing of manuscript)

## Acknowledgements

The authors thank Drs. Jonathan Livingston, Dwayne Brandon and Sandra Waters from the Department of Psychology, North Carolina Central University, for helpful discussions and suggestions during the design and analysis of the study. The authors also thank Mr. Lavonza Holliman for his assistance with the initial literature review.

